# Symmetry breaking in the female germline cyst

**DOI:** 10.1101/2021.05.07.443143

**Authors:** D. Nashchekin, L. Busby, M. Jakobs, I. Squires, D. St Johnston

**Author notes:** Institute for Cell and Molecular Biosciences, Newcastle University, Newcastle upon Tyne NE2 4HH, UK.

## Abstract

In mammals and flies, only a limited number of cells in a multicellular female germline cyst become oocytes, but how the oocyte is selected is unknown. Here we show that the microtubule minus end-stabilizing protein, Patronin/CAMSAP marks the future *Drosophila* oocyte and is required for oocyte specification. The spectraplakin, Shot, recruits Patronin to the fusome, a branched structure extending into all cyst cells. Patronin stabilizes more microtubules in the cell with most fusome and this weak asymmetry is amplified by Dynein-dependent transport of Patronin-stabilized microtubules. This forms a polarized microtubule network, along which Dynein transports oocyte determinants into the presumptive oocyte. Thus, Patronin amplifies a weak fusome anisotropy to break cyst symmetry. These findings reveal a molecular mechanism of oocyte selection in the germline cyst.

---

In many organisms, only a subset of female germ cells develop into oocytes, with the rest becoming accessory cells that contribute cytoplasm and organelles to the oocytes (*1*). For example, mouse female germ cells form cysts of up to 30 cells, but most cells undergo apoptosis after transferring cytoplasm and centrosomes to the small number of cells that become oocytes (*2, 3*). In *Drosophila*, germline cyst formation starts in the germarium at the anterior tip of the ovary, when a stem cell divides asymmetrically to produce another stem cell and a cystoblast, which then divides four times with incomplete cytokinesis to generate a cyst of 16 germ cells connected by intercellular bridges called “ring canals” (*4, 5*). As the cyst moves posteriorly through region 2a-b of the germarium, it is surrounded by epithelial follicle cells that separate it from other cysts. The cyst then rounds up in region 3 to form a follicle. By this stage, one cell has been selected as the oocyte, while the other 15 become nurse cells (Fig. 1A). Oocyte selection depends on the formation of a noncentrosomal microtubule organizing centre (ncMTOC) in the future oocyte that organizes a polarized microtubule network that directs the dynein-dependent transport of cell fate determinants and centrosomes from the other 15 cyst cells into the future oocyte (*6*-*8*) (Fig. 1A). How symmetry is broken to specify which cell contains the ncMTOC and becomes the oocyte is unclear.

**Fig. 1.**
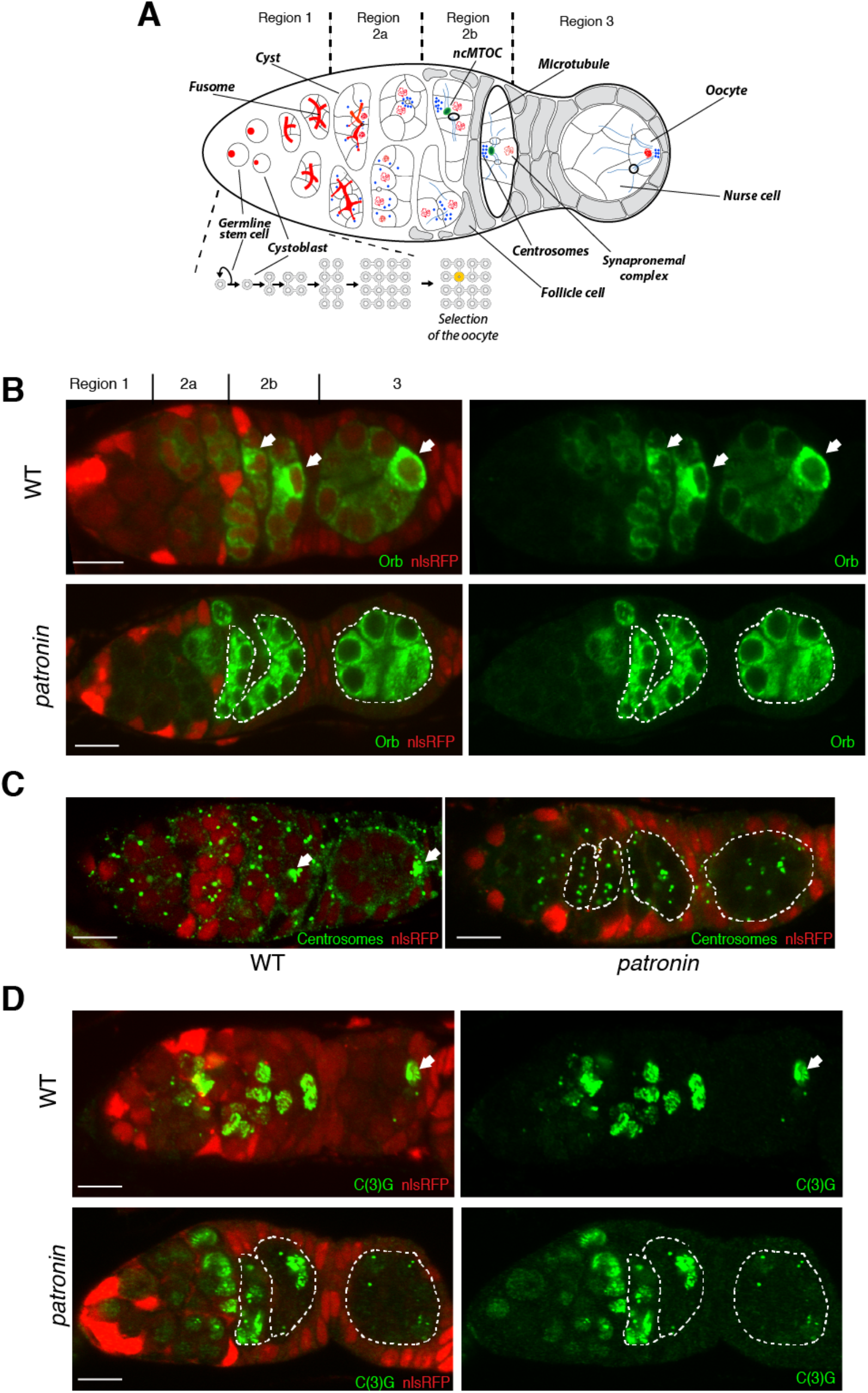
Patronin is required for the oocyte specification. **(A)** A schematic diagram of *Drosophila* germarium showing germline cyst formation and oocyte selection. **(B-D)** Distribution of oocyte specification markers Orb **(B)**, centrosomes **(C)** and C(3)G **(D)** in wild type (WT; top or left in C) and *patronin* mutant (bottom or right in C) cysts. Arrows point to the future oocyte. Mutant cysts are labeled by the absence of nlsRFP and are marked by dashed lines. Regions of the germarium are indicated on the top. Scale bar, 10µm.

Patronin and its vertebrate orthologues, the CAMSAPs are microtubule minus end binding proteins that have been recently found to be essential components of ncMTOCs (*9*-*13*). To investigate the role of Patronin in oocyte determination, we examined the distribution of oocyte markers in *patronin*^*c9-c5*^ mutant germline cysts (Fig. 1B-D). In wild-type cysts, Orb and centrosomes accumulate in the future oocyte in region 2b and 3 (*14*-*16*), but they fail to localise in *patronin* mutants and remain at similar levels in all 16 cells of the cyst (76% and 97% of mutant cysts respectively) (Fig. 1B-C). Several germ cells normally enter meiosis in region 2a and accumulate the synaptonemal complex protein C(3)G. C(3)G then becomes restricted to two cells in region 2b and to just the oocyte in region 3 (*17*) (Fig. 1D). By contrast, C(3)G is not localised in region 3 of *patronin* cysts and 44% of the cysts in region 2b have 3 cells in meiosis (Fig. 1D). These phenotypes show that Patronin is required for oocyte determination.

To examine whether Patronin is asymmetrically distributed in the cyst we imaged live germaria expressing endogenously tagged Patronin-Kate. Patronin starts to accumulate in a single cell in each cyst in region 2a, which is earlier than other markers for the presumptive oocyte and remains in one cell in regions 2b and 3, where it forms distinct foci in the cytoplasm (Fig. 2A-2A’). This cell will become the oocyte, as it is also labelled by Orb (Fig. 2B) and C(3)G (Fig. 2C). *patronin* mRNA is not localised within the cyst and Patronin expressed from a cDNA with heterologous UTRs under the control of the *ubiquitin* promoter shows a similar distribution to an endogenous protein, indicating that Patronin is localised as a protein and not through transcription in this cell or mRNA localisation (Fig. 2B-C and Fig. S1).

**Fig. 2.**
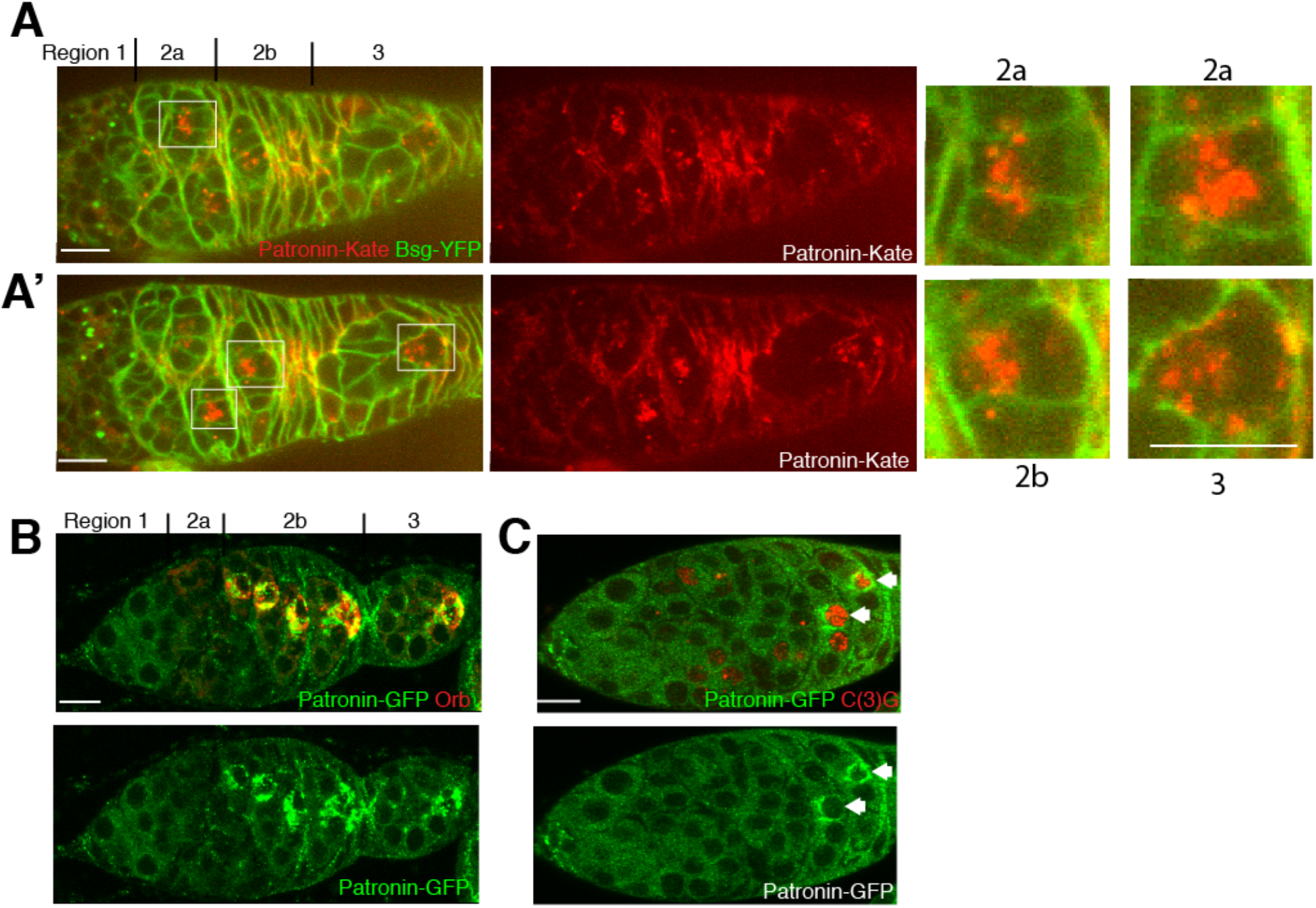
Patronin accumulates in the future oocyte. **(A-A’)** Two different focal planes of a live germarium showing accumulation of endogenously tagged Patronin-Kate in one cell of the cyst. Regions 2a, 2b and 3 are shown as close-ups. Cell membranes are labelled by Basigin-YFP (Bsg-YFP). **(B-C)** Ectopically expressed Patronin-GFP accumulates in future oocytes labelled by Orb **(B)** or C(3)G **(C).** Arrows point to the future oocyte. Regions of the germarium are indicated on the top. Scale bar, 10µm.

Like Orb, Dynein does not localise to the presumptive oocyte in *patronin* mutant cysts (Fig. S2). This suggests that the loss of Patronin disrupts the formation of the MTOC in the future oocyte, leading to loss of the polarised microtubule network along which Dynein transports cargoes into one cell. As most of MT plus ends accumulate at the site of MT nucleation, we used the MT plus end tracking protein EB1-GFP to visualise the putative MTOC in the cyst. In wild-type cysts, the majority of EB1-GFP comets localise to one cell in region 2b and 3 (Fig. 3A-B and Movie S1 and S2). Moreover, the densest EB1-GFP signal co-localises with the Patronin foci in the same cell, suggesting that the latter are the MTOCs formed in the future oocyte (Fig. 3C). This asymmetric distribution of EB1-GFP is lost in *patronin* mutant cysts, where EB1-GFP comets are distributed more homogeneously (Fig. 3A-B and Movie S3 and S4). Patronin is therefore required for MTOC formation in the presumptive oocyte and the formation of a polarised MT network.

**Fig. 3.**
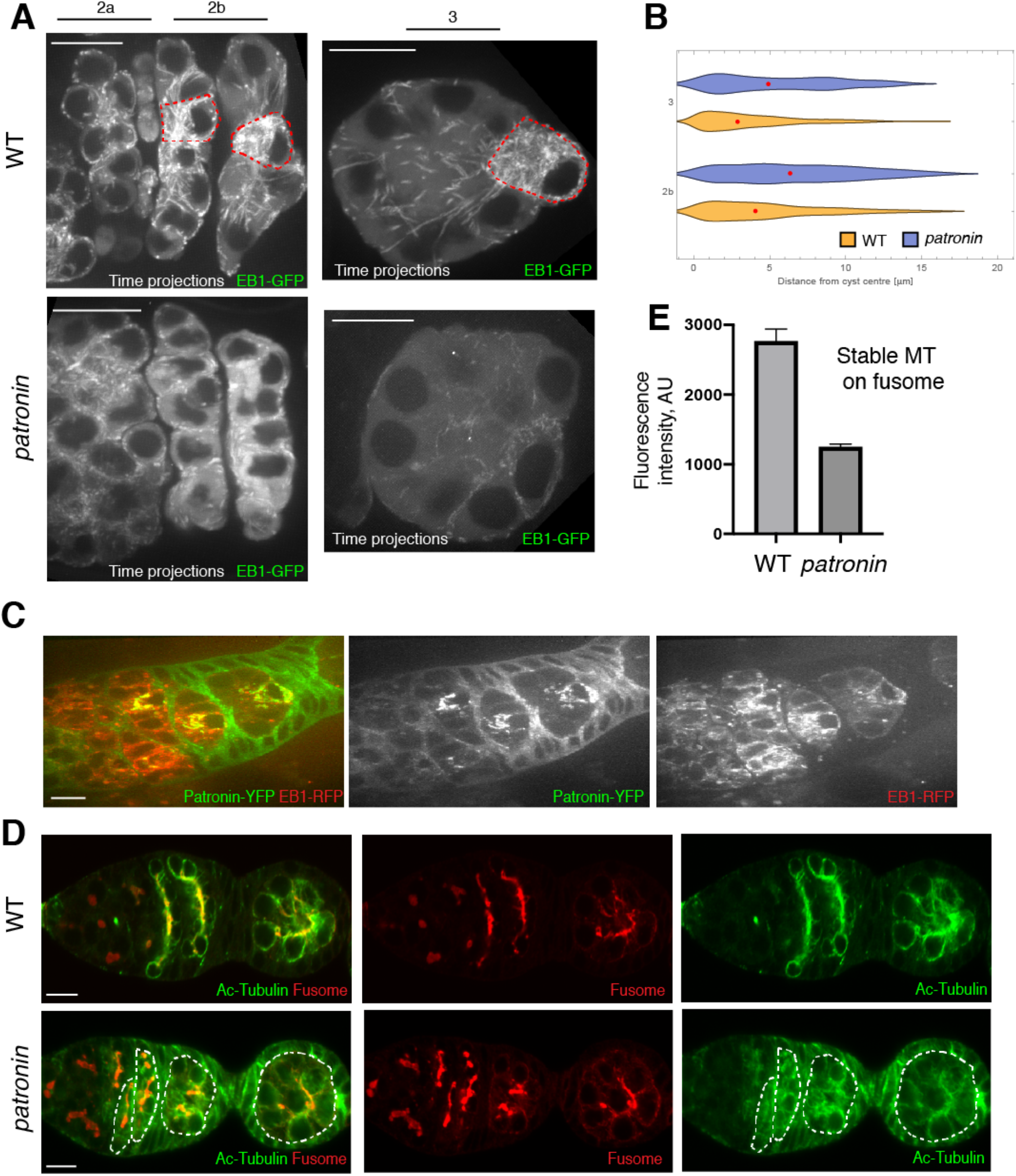
Patronin is required for MT organisation in the cyst. **(A-C)** Patronin is required for MTOC formation in the presumptive oocyte. **(A)** EB-1 comet tracks in wild type (WT; top) and *patronin* mutant (bottom) cysts. The images are projections of several time points from Movies S1 (WT; region 2), S2 (WT; region 3), S3 (*patronin*; region 2), S4 (*patronin*; region 3). The red dashed line marks cells with MTOCs. Regions of the germarium are indicated on the top. **(B)** Quantification of EB-1 comets distribution in wild type (WT) and *patronin* mutant cysts in region 3 and 2b of germarium. Red dots indicate median value. **(C)** Live germarium shows co-localisation of Patronin-YFP foci with microtubules plus end marker EB1-GFP in the presumptive oocyte. **(D-E)** Patronin stabilises microtubules in the cyst. **(D)** Wild type (WT; top) and *patronin* mutant (bottom) cysts stained with anti-acetylated tubulin (Ac-Tubulin) and anti-Shot (Fusome). Mutant cysts are marked by dashed lines. **(E)** Quantification of a mean fluorescence intensity of fusome associated acetylated microtubules in *patronin* mutant and WT cysts. Errors bars indicate the SEM. Scale bar, 10µm.

Wild-type cysts contain a population of stable, acetylated MTs that form along the fusome, an ER, spectrin and actin-rich structure that connects all 16 cells of the cyst (*16*-*19*) (Fig. 3D). In *patronin* mutant cysts, there is a 2.5 fold reduction in stable MTs (Fig. 3D-E). Thus, in the absence of Patronin the whole organisation of MTs in the cyst is disrupted. Patronin binds MT minus ends and stabilises MTs by protecting theirs minus ends against kinesin-13 induced depolymerisation (*11, 13*). Our results indicate that early accumulation of Patronin in only one cell of the cyst stabilises MT minus ends there, leading to dynein-dependent transport into this cell, the formation of MTOCs and subsequent specification as the oocyte.

To examine whether centrosomes contribute to the formation of Patronin MTOCs, we imaged germline cysts expressing endogenously tagged Patronin-YFP and the centrosomal protein Asterless fused to Cherry. Although centrosomal clusters localise near Patronin foci, the Asterless and Patronin signals only partially overlap and most Patronin foci lie outside the centrosomal cluster, indicating that Patronin MTOCs are noncentrosomal (Fig. S3A). Centrosomes have been proposed to be inactive during their migration into the oocyte and they lack crucial components of the PCM(*8*). To test whether centrosomes contribute to microtubule organisation, we imaged germline cysts expressing EB1-GFP and Asterless-Cherry. The centrosomes show strong MT nucleating activity in region 1, where they organise the mitotic spindles (Fig. S3B and Movie S5). However, only some Asterless-Cherry labelled centrosomes in the presumptive oocyte produce EB1-GFP comets in region 2b (Fig. S3C and Movie S6). Thus, Patronin-dependent ncMTOCs create the initial asymmetry in MT organisation that leads to the accumulation of centrosomes in the future oocyte, which may then be amplified by activation of some centrosomes in this cell. The close proximity of centrosomal clusters to ncMTOCs, raises the possibility that new MTs produced by centrosomes are released and then captured and stabilised by Patronin in ncMTOCs, a mechanism described for CAMSAP proteins (*20*). Partial centrosomal activity could also explain the presence of EB-GFP comets in *patronin* mutant cysts (Fig. 3A).

The observation that Patronin is the earliest marker for the future oocyte raises the question of how symmetry is broken in the cyst to enrich Patronin in one cell. One proposed mechanism for symmetry-breaking is that the cell that inherits the most fusome becomes the presumptive oocyte (*21*). The fusome is asymmetrically partitioned during the mitoses in region 1, so that mother cells inherit more material than their daughters and one of the two cells with four ring canals has more fusome than the rest (*19*). To examine whether Patronin associates with the fusome, we imaged live germaria expressing endogenously-tagged Patronin-YFP and the fusome marker, Hts-Cherry (Hu li tai shao). Patronin localizes on the fusome in early region 2a, but becomes concentrated in one cell as the cyst progresses towards region 3 (Fig. 4A). When the MTs are depolymerized with colcemid, however, Patronin remains on the fusome in regions 2b and 3 (Fig. 4B). Thus, the fusome determines the initial localisation of Patronin in early region 2a, including its slight enrichment in the future oocyte, which is then amplified by a MT-dependent process.

**Fig. 4.**
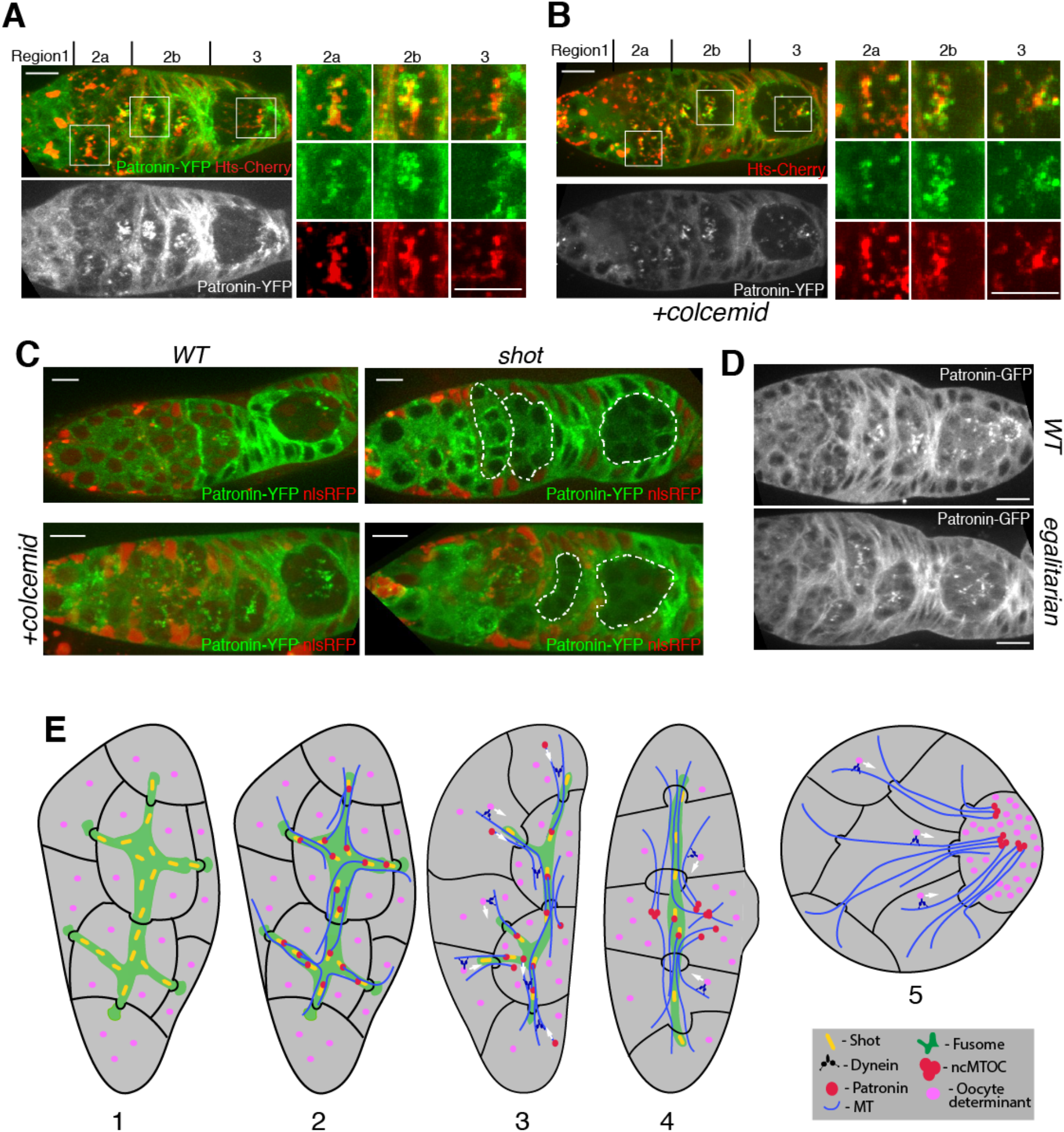
Patronin localisation is defined by fusome and by a positive feed back loop of Dynein mediated transport. **(A-B)** Patronin associates with fusome in microtubules-dependent manner. Live germaria untreated **(A)** or treated with colcemid **(B)** expressing Patronin-YFP and a fusomal protein Hts-Cherry. Regions 2a, 2b and 3 are shown as close-ups. Regions of the germarium are indicated on the top. **(C)** Shot links Patronin to the fusome. Live germaria with wild type (WT; left) and *shot* mutant (right) cysts untreated (top) and treated with colcemid (bottom) expressing Patronin-YFP. Mutant cysts are labelled by the absence of nlsRFP and are marked by dashed lines. **(D)** Patronin localisation depends on Dynein activity. Wild type (WT; top) and *egalitarian* mutant (bottom) live germaria expressing transgenic Patronin-GFP. **(E)** A 4-step model of cyst polarisation (1-4). A region 3 cyst with the oocyte positioned at the posterior of the cyst (5). See text for details. Scale bar, 10µm.

The spectraplakin, Shot, localises to the fusome, is required for the oocyte specification and recruits Patronin to ncMTOCs in the oocyte later in oogenesis, making it a good candidate for a factor that links Patronin to the fusome (*13, 17*). In *shot* mutant cysts, Patronin does not accumulate in one cell of the cyst and fails to form foci (Fig. 4C). Furthermore, loss of Shot prevents Patronin from associating with the fusome when the MTs are depolymerised (Fig. 4C). Thus, Shot is required to recruit Patronin to the fusome, thereby transmitting fusome asymmetry to Patronin localisation.

The MT-dependent enrichment of Patronin in one cell as the cyst moves through the germarium suggests its initial, weakly asymmetric distribution on the fusome is then amplified by Dynein-dependent transport towards the minus ends of the MT that have been stabilized by Patronin. Dynein mutants disrupt fusome formation and the early divisions of the cyst (*7*). We therefore tested its function by examining components of the Dynein/dynactin complex that are specifically required for oocyte specification: *egl, BicD* and *Arp1* (*22*-*24*), (Fig. 4D, S4A and S4B). Like MT depolymerisation, mutations in any of these genes disrupt the enrichment of Patronin foci in one cell. Deletion of the MT minus end binding domain of Patronin, but not the CKK domain (*25*), also prevents Patronin accumulation in the pro-oocyte (Fig. S4C-D). Thus, Patronin localisation depends on its binding to MT minus ends and on Dynein activity, suggesting that Dynein transports Patronin bound to MT minus ends towards the future oocyte.

Our observations lead us to propose a 4-step model of cyst polarisation and oocyte selection (Fig. 4E). 1) During germ cell divisions and cyst formation, the asymmetric segregation of the fusome leads to the one cell with more fusome material than the rest. 2) In region 2a, Patronin is recruited to the fusome by Shot. The cell with most fusome therefore contains more Patronin, leading to the stabilisation of more MT minus ends in this cell and a weakly polarised MT network. 3) Both Patronin bound MTs in other cells of the cyst and oocyte determinants are then transported by Dynein along these MTs towards their minus ends in the presumptive oocyte. 4) This creates a positive feedback loop: as Dynein transports more Patronin and MTs into the cell with most stabilised MT minus ends, more minus ends become stabilised in this cell, amplifying the MT polarity and leading to enhanced Dynein transport of its cargoes into this cell. In this way, the small original asymmetry in the fusome is converted into the highly polarised MT network that concentrates the oocyte determinants in one cell.

The selection of the oocyte from a cyst of 16 germ cells is the first symmetry-breaking step in *Drosophila* oogenesis and is an essential prerequisite for the establishment of the anterior-posterior axis, which occurs in region 3 of the germarium, when the oocyte is positioned at the posterior of the cyst (*26*). We identify Patronin as one of the earliest factors that defines the future oocyte and Patronin ncMTOC as a key structure in organising the polarised MT network in the cyst. Patronin is a member of the conserved CAMSAP family of minus end binding proteins, raising the possibility that the molecular mechanisms of oocyte selection in *Drosophila* could be conserved during the formation of mammalian oocytes. Although fusomes have not been observed in mammalian cysts (*27*), MT-dependent transport of organelles through intercellular bridges has been shown to play an important role in oocyte differentiation in mice (*3*).

## Materials and Methods

### Mutant alleles

The following *Drosophila melanogaster* mutant alleles have been described previously and can be found on FlyBase.org: *shot*^*3*^ (*28*), *BicD*^*r5*^ (*29*), *arp1*^*c04425*^ (*30*), *egalitarian*^*1*^ *and egalitarian*^*2*^ (*31*). The *patronin*^c9-c5^ was generated by injecting nos::Cas9 embryos(*32*) with a sgRNA targeting the second protein coding exon in *patronin* (5’ GGACAT**GC**CCATTACC**G**AAA 3’). *patronin*^c9-c5^ has a substitution of GC (highlighted in bold in sgRNA sequence) to TA changing Met Pro to Ile Ser; and deletion of G (highlighted in bold in sgRNA sequence) changing Glu Thr Val Leu to Lys Arg Tyr STOP.

### Fluorescent marker stocks

Hts-Cherry was derived by N. Lowe from the Hts-GFP CPTI protein trap line using P-element exchange (*33, 34*). Basigin-YFP is Cambridge Protein Trap Insertion line (*33*). The following stocks have been described previously: Asterless-Cherry (*35*), UAS EB1-GFP (*36*), Patronin-YFP (*13*), pUbq-Patronin-GFP (*37*) (A isoform). Patronin-mKate, UAS EB1-RFP, pUbq-Patronin-GFP (I isoform), pUbq-PatroninΔMTD-GFP, pUbq-PatroninΔCKK-GFP were from this study.

### *Drosophila* genetics

Germline clones of *patronin*^c9-c5^, *shot*^*3*^, *BicD*^*5*^, *arp1*^*c04425*^ were induced by incubating larvae at 37° for two hours per day over a period of three days. Clones were generated with FRT G13 nlsRFP, FRT 40A nlsRFP, FRT 82B nlsRFP (Bloomington Stock Center) using the heat shock Flp/FRT system (*38*). Germline expression of UAS EB1-GFP and UAS EB1-RFP was induced by nanos-Gal4.

### Molecular Biology

The Patronin C-terminal mKate knockin was made by injecting nos::Cas9 embryos (*32*) with a single guide RNA targeting the region of the stop codon in *patronin* (5’-GGCGCTTGTAATCTAAGCGG-3’) and a donor plasmid with 4-kb homology arms surrounding the mKate sequence. A full-length *patronin* RI cDNA was amplified from pUASP mCherry-Patronin (*13*) and cloned together with EGFP into pUbq-attb vector downstream of the polyubiqutin promoter. pUbq-PatroninΔMTD-GFP and pUbq-PatrioninΔCKK-GFP were generated by PCR amplifying the corresponding fragments (see Ref (25) for details) from pUbq-Patronin-GFP and cloning them into pUbq-attb. EB1 cDNA was amplified from pUASP EB1-GFP and cloned together with tagRFP into pUASP-attb to generate UAS EB1-RFP.

### Immunohistochemistry

Ovaries were fixed for 20 min in 4% paraformaldehyde and 0.2% Tween in PBS. Ovaries were then blocked with 1% BSA in PBS for 1 hr at room temperature. Ovaries were incubated with the primary antibody for 16 hr with 0.1% BSA in PBS with 0.2% Tween at 4C and for 4 hr with the secondary antibody at room temperature. We used the following primary antibodies: mouse anti-acetylated tubulin at 1:250 (Sigma); mouse anti-Dynein heavy chain at 1:50 (DSHB Hybridoma Product 2C11-2. Deposited to the DSHB by Scholey, JM); guinea pig anti-Shot (*13*) at 1:500, rabbit anti-dPLP at 1:1000 (gift from J. Raff, University of Oxford, UK), mouse anti-C(3)G at 1:500 (gift from R.S. Hawley, Stowers Institute, US), mouse anti-Orb at 1:10 (DSHB Hybridoma Products 4H8 and 6H4. Deposited to the DSHB by Schedl, P). Conjugated secondary antibodies (Jackson Immunoresearch) were used at 1:100. In situ hybridizations were performed as previously described (*24*).

### Colcemid Treatment

Flies were starved for 2 hr and then fed colcemid (Sigma) in yeast paste (66 µg/ml) for 17 hr. Ovaries were dissected and imaged as described below.

### Imaging

For live imaging, ovaries were dissected and imaged in Voltalef oil 10S (VWR International) on an Olympus IX81 inverted microscope with a Yokogawa CSU22 spinning disk confocal imaging system (60x/ 1.35 NA Oil UPlanSApo and 100x/ 1.3 NA Oil UPlanSApo) or on Leica SP5 confocal microscope (63x/1.4 HCX PL Apo CS Oil). To label cell membranes, ovaries were dissected in Schneider’s medium (Sigma) with 10 µg/ml insulin (Sigma) and CellMask (1:2000, Life Technologies), incubated for 10 min at room temperature, washed and transferred to Voltalef oil for imaging. Fixed preparations were imaged using an Olympus IX81 (60x/ 1.35 NA Oil UPlanSApo) or a Leica SP8 (63x/1.4 HC PL Apo CS Oil) confocal microscope. Images were collected with Olympus Fluoview, MetaMorph and Leica LAS AF software and processed using ImageJ. Germaria were imaged by collecting 10–15 z sections spaced 0.5 µm apart. The images in Fig. 2B, Fig. 4A, Fig. 4B, Fig. 4C (bottom panels) and Fig. 4D are projections of several z sections.

### EB1 tracking

EB1 comets were tracked using the ImageJ plugin TrackMate. For each movie, tracking performance was visually inspected and optimal tracking parameters chosen accordingly. To determine the distribution of EB1 comets in the cyst we calculated the distance from each EB1 track starting point to the cyst centre. The following number of comets were analysed: WT region 3: 910 comets, WT region 2b: 430 comets; *patronin* mutant region 3: 989 comets, *patronin* mutant region 2b: 1736 comets.

### Statistical analyses

The chi-square test was used to test whether values were significantly different between WT and *patronin* mutant cysts. The Mann-Whitney t-test was used to determine significance when comparing fluorescence intensity of acetylated tubulin staining. We used the MATLAB implementation of the Kruskal Wallis Test followed by a Tukey post hoc test to determine statistical differences in EB-1 comet distributions. No statistical methods were used to predetermine sample size, the experiments were not randomized, and the investigators were not blinded to allocation during experiments and outcome assessment.

### Reproducibility of experiments

Images are representative examples from at least three independent repeats for each experiment. The number of cysts analyzed for each experiment were as follows: Fig. 1B (WT 27, *patronin* 27), Fig, 1C (WT 30, *patronin* 41), Fig. 1D (WT 33, *patronin* 31), Fig. 2A (17), Fig. 2B (53), Fig. 2C (32), Fig. 3A (WT 30, *patronin* 29), Fig. 3C (41), Fig3D (WT 40, *patronin* 45), Fig. 3E (WT 18, *patronin* 6), Fig. 4A (30), Fig. 4B (54), Fig. 4C (WT 35, WT + colcemid 37, *shot* 28, *shot* + colcemid 31), Fig. 4D (WT 40, *egalitarian* 34).

## Acknowledgments

We are grateful to R. Hawley, J. Raff, J. Scholey and the Bloomington Stock Center (NIH P40OD018537) for flies and reagents, the Gurdon Institute Imaging Facility for assistance with microscopy. We thank N. Lowe for technical assistance.

## Funding

Wellcome Principal Research Fellowship 080007 and 207496 (DSJ)

Wellcome core support 092096 and 203144 (DSJ)

Wellcome PhD studentship 109145 (MJ)

Cancer Research UK core support A14492 and A24823 (DSJ)

BBSRC grant BB/R001618/1 (DSJ, DN, IS)

## Author contributions

Conceptualization: DN, DSJ

Methodology: DN, MJ

Investigation: DN, LB, MJ, IS, DSJ

Visualization: DN, MJ, DSJ

Funding acquisition: DN, DSJ

Project administration: DN, DSJ

Supervision: DN, DSJ

Writing – original draft: DN, DSJ

Writing – review & editing: DN, LB, MJ, IS, DSJ

## Data and materials availability

All data are available in the main text or the supplementary materials.

---

**Fig. S1.**
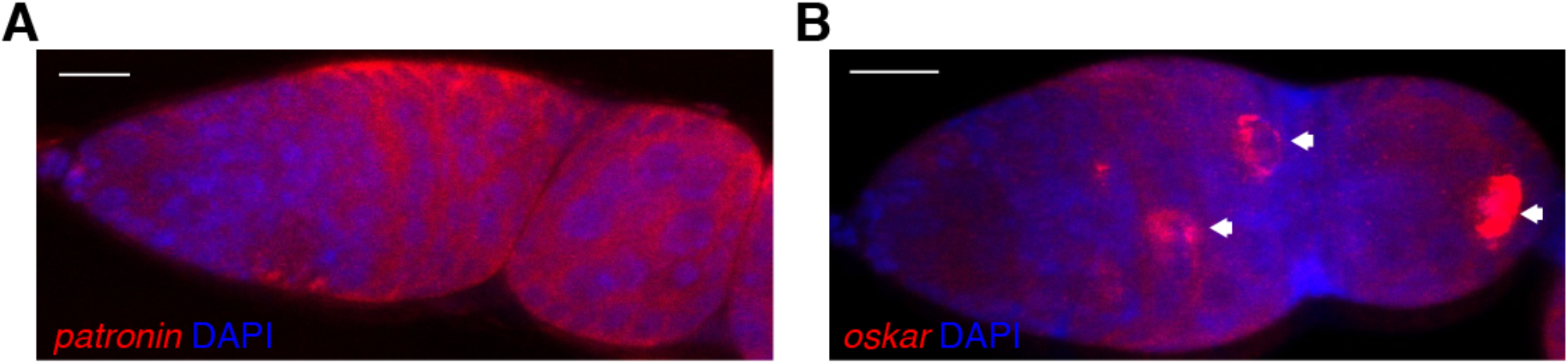
Patronin mRNA is not localised in the cyst. Confocal images of fluorescent in situ hybridisations (FISH) to endogenous *patronin* **(A)** and *oskar* **(B)** mRNA in a wild type germarium, counterstained with DAPI to label the nuclei. Arrows point to the future oocyte. Scale bar, 10µm.

**Fig. S2.**
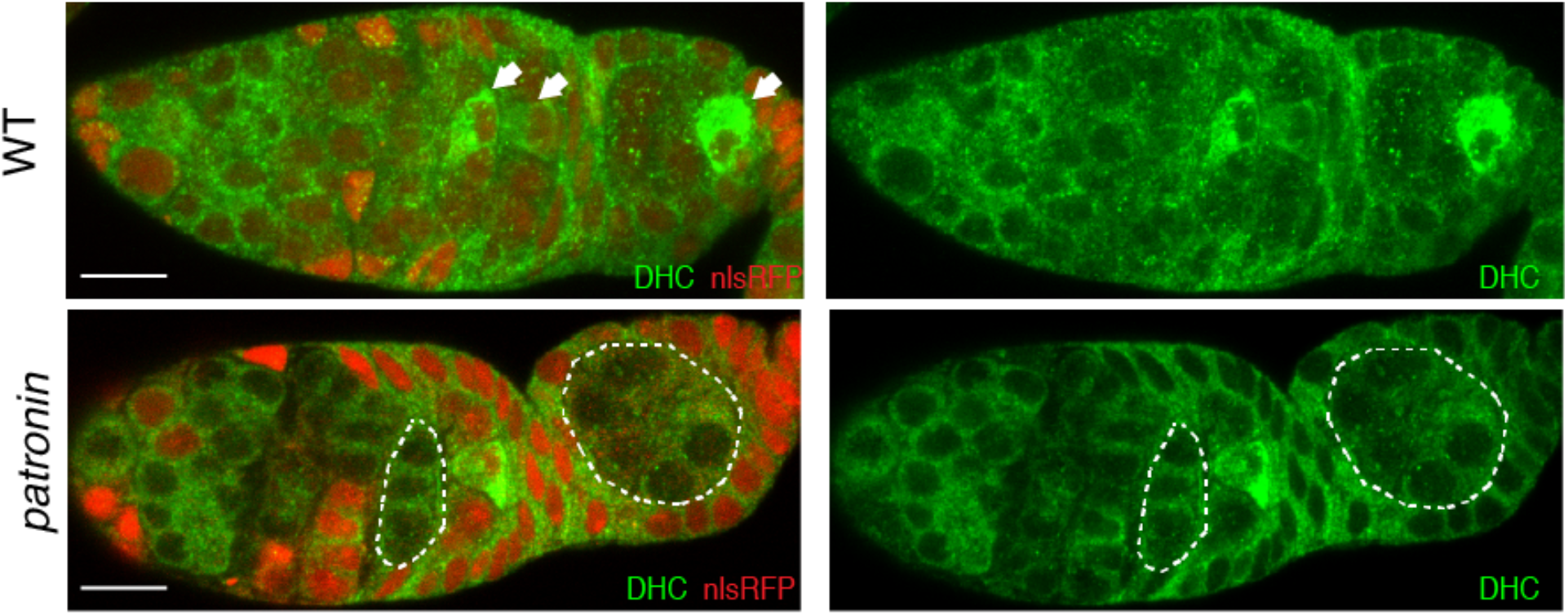
Patronin is required for Dynein localisation. Distribution of Dynein Heavy Chain (DHC) in wild type (WT; top) and *patronin* mutant (bottom) cysts. Arrows point to the future oocyte. Mutant cysts are labeled by the absence of nlsRFP and are marked by the dashed line. Scale bar, 10µm.

**Fig. S3.**
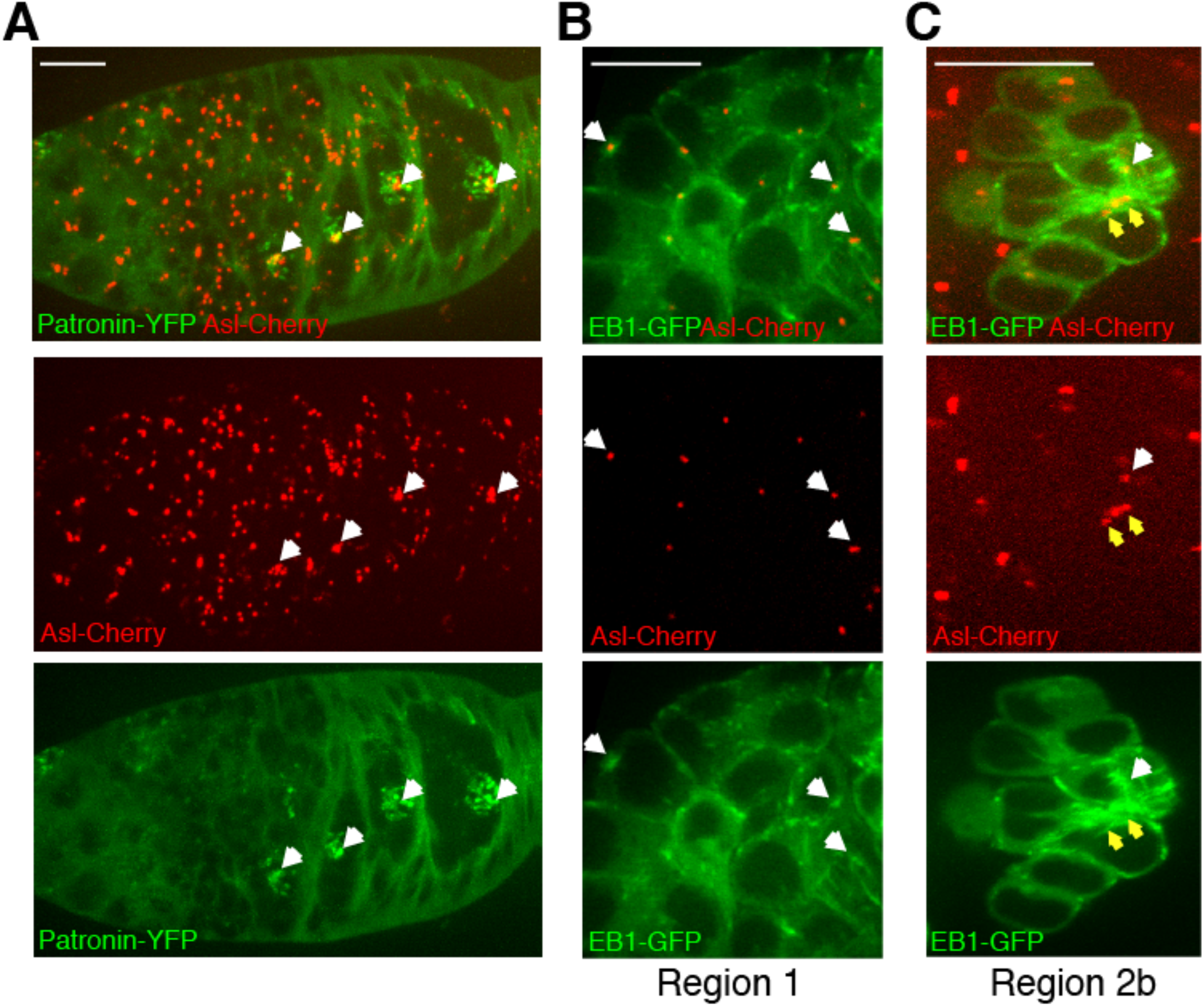
Patronin MTOCs are not centrosomal. **(A)** Patronin foci lie outside the centrosomal cluster. Live germarium expressing Patronin-YFP and a centrosomal protein Asterless-Cherry (Asl-Cherry). The image is a projection of several z sections spanning the cyst. Arrows mark centrosomal clusters. **(B-C)** Centrosomes contribute to microtubule organisation in the cyst. Live germaria expressing EB1-GFP and Asterless-Cherry (Asl-Cherry). **(B)** Region 1 of the germarium. The image is taken from Movie S5. Arrows indicate active centrosomes. **(C)** Region 2b of the germarium. The images are projections of several time points from Movie S6. The white arrow points to an active centrosome in the presumptive oocyte. Yellow arrows point to two inactive centrosomes. Scale bar, 10µm.

**Fig. S4.**
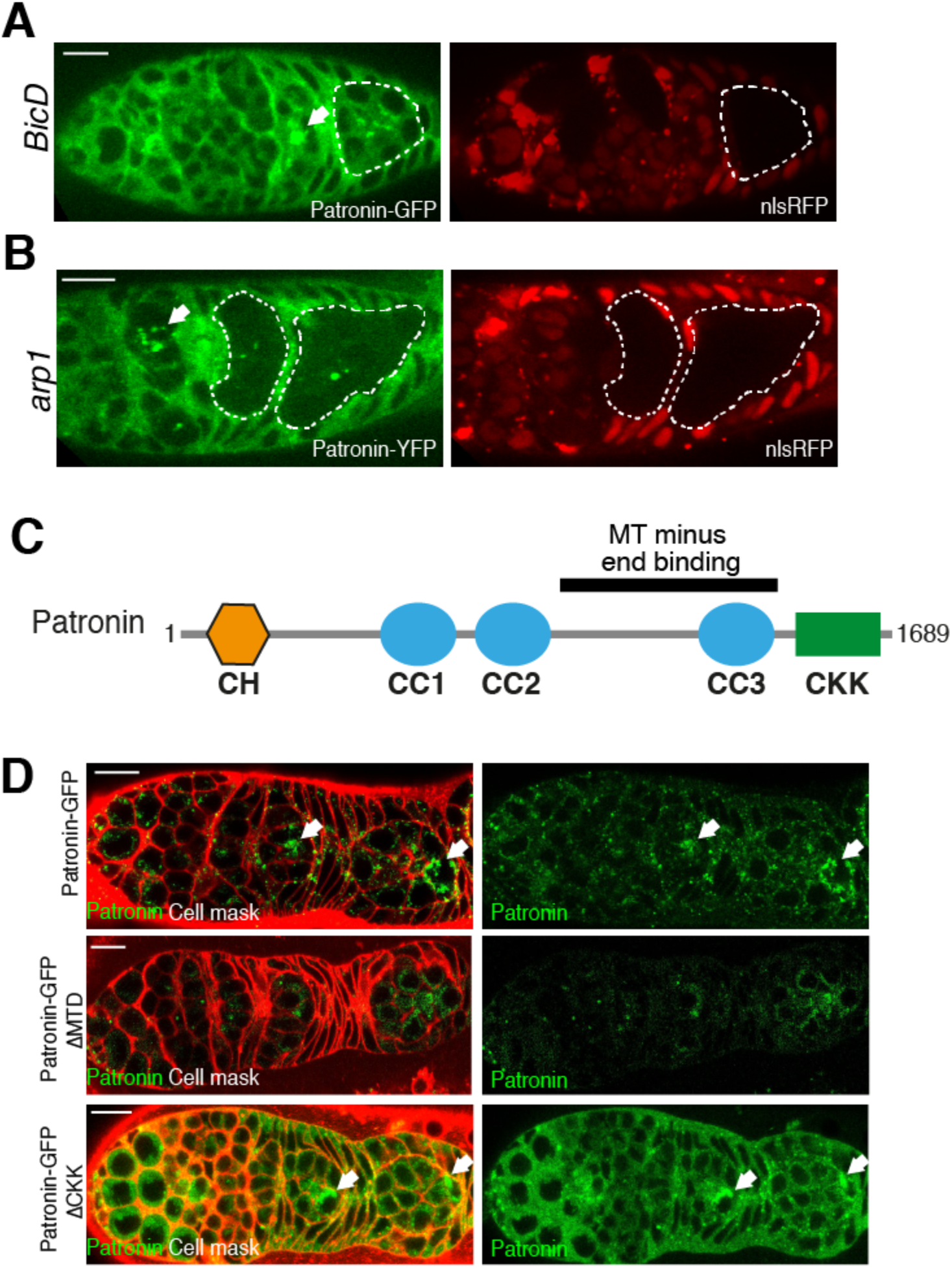
Patronin localisation depends on Dynein activity and MT minus ends binding. **(A-B)** Live germaria with *BicD* **(A)** or *arp1* **(B)** mutant cysts expressing transgenic Patronin-GFP **(A)** or Patronin-YFP **(B)**.Mutant cysts are labeled by the absence of nlsRFP and are marked by a dashed line. Arrows indicate wild type cysts. **(C)** A diagram of Patronin domain structure. **(D)** MT minus end binding domain (MTD) is required for Patronin localisation. Live germaria expressing wild type (top), MTD-deleted (middle) and CKK-deleted (bottom) transgenic Patronin-GFP. Cell membranes are labelled by CellMask. Arrows indicate accumulation of Patronin foci in the presumptive oocyte. Scale bar, 10µm.

**Movie S1**. A time-lapse video of the microtubule plus-end binding protein EB1-GFP in wild-type germline cysts in region 2 of the germaium. The red line outlines the cells containing MTOCs. Related to Figure 3A. Images were collected every 1 second on a spinning disc confocal microscope. The video is shown at 15 frames/sec.

**Movie S2**. A time-lapse video of the microtubule plus-end binding protein EB1-GFP in wild-type germline cysts in region 3. Related to Figure 3A. Images were collected every 1 second on a spinning disc confocal microscope. The video is shown at 15 frames/sec.

**Movie S3**. A time-lapse video of the microtubule plus-end binding protein EB1-GFP in *patronin* mutant germline cysts in region 2. Related to Figure 3A. Images were collected every 1 second on a spinning disc confocal microscope. The video is shown at 15 frames/sec.

**Movie S4**. A time-lapse video of the microtubule plus-end binding protein EB1-GFP in *patronin* mutant germline cysts in region 3. Related to Figure 3A. Images were collected every 1 second on a spinning disc confocal microscope. The video is shown at 15 frames/sec.

**Movie S5**. A time-lapse video of the microtubule plus-end binding protein EB1-GFP (green) and the centrosomal protein Asterless-Cherry (red) in wild-type germline cysts in region 1. Arrowheads point to active centrosomes. Related to Figure S3B. Images were collected every 1.6 seconds on a spinning disc confocal microscope. The video is shown at 15 frames/sec.

**Movie S6**. A time-lapse video of the microtubule plus-end binding protein EB1-GFP (green) and centrosomal protein Asterless-Cherry (red) in wild-type germline cysts in region 2b. The white arrow points to an active centrosome in the presumptive oocyte. Related to Figure S3C. Images were collected every 1.6 seconds on a spinning disc confocal microscope. The video is shown at 15 frames/sec.

